# Independent Control of the Thermodynamic and Kinetic Properties of Aptamer Switches

**DOI:** 10.1101/688275

**Authors:** Brandon D. Wilson, Amani A. Hariri, Ian A.P. Thompson, Michael Eisenstein, H. Tom Soh

**Author notes:** equal contribution (Brandon D. Wilson, Amani A. Hariri).

## Abstract

Molecular switches that change their conformation upon target binding offer powerful capabilities for biotechnology and synthetic biology. In particular, aptamers have proven useful as molecular switches because they offer excellent binding properties, undergo reversible folding, and can be readily engineered into a wide range of nanostructures. Unfortunately, the thermodynamic and kinetic properties of the aptamer switches developed to date are intrinsically coupled, such that high temporal resolution (*i.e.*, switching time) can only be achieved at the cost of lower sensitivity or high background. Here, we describe a general design strategy that decouples the thermodynamic and kinetic behavior of aptamer switches to achieve independent control of sensitivity and temporal resolution. We used this strategy to generate an array of aptamer switches with effective dissociation constants (*K*_*D*_) ranging from 10 μM to 40 mM and binding kinetics ranging from 170 ms to 3 s—all generated from the same parent ATP aptamer. Our strategy is broadly applicable to other aptamers, enabling the efficient development of switches with characteristics suitable for diverse range of biotechnology applications.

## Introduction

A wide range of essential biological functions are governed by the action of molecular ‘switches’^1,2^, which undergo a reversible conformational change upon binding a specific target molecule. These switches can be coupled to other molecular machinery to trigger a wide range of downstream functions. There is considerable interest in engineering biologically inspired molecular switches that can achieve a selective and sensitive output in response to binding a target molecule, which could prove valuable for diverse applications, including imaging^3,4^, biosensing^5^, and drug delivery^6,7^. Aptamers have proven to be particularly promising and versatile in this regard^8^ as they are highly stable, easy to synthesize, exhibit reversible binding, and are readily adaptable to chemical modification. Since conventional methods of aptamer generation do not routinely yield aptamers capable of structure switching, many selection schemes^9,10^ and engineering approaches^11–13^ have been developed for the creation of aptamer switches. In contrast to naturally occurring molecular switches that have evolved over millions of years to function under precise physiological conditions, switches based on synthetic affinity reagents must be tuned to match their intended function. Unfortunately, existing selection and engineering strategies offer limited control over the thermodynamic and kinetic properties of the resultant aptamer switches and, by extension, over properties such as effective binding affinity and temporal resolution.

Recent work has yielded important insights into how to control the binding of molecular switches. For instance, it has been shown that the hybridization strength of the hairpin in a molecular beacon can modulate the effective detection range for target concentrations spanning many orders of magnitude^14^. Recognizing that this enthalpy-driven control is coarse-grained, other work achieved fine-grained control of effective binding affinity through the modulation of the entropic change associated with the degree of confinement imposed by a intramolecular linker of variable length^15^. However, previous efforts at tuning the binding properties of aptamer constructs have found that their thermodynamic and kinetic properties are intrinsically coupled^16^, such that fast temporal resolution can only be achieved at the cost of either large background signal or lower affinity, or requires high-temperature conditions that can interfere with ligand binding^17^. At present, there is no reliable strategy for independently controlling the thermodynamic and kinetic properties of engineered aptamer switches.

Here, we introduce a general framework for the design of aptamer switches that enables independent control over their thermodynamic and kinetic properties. We explore the use of an intramolecular strand-displacement (ISD) strategy^18^ and the degree to which the binding properties of this construct can be controlled through rational design. Our ISD construct consists of a single-molecule switch in which an aptamer is attached to a partially complementary displacement strand via a poly-T linker. Briefly, target binding to the aptamer shifts the equilibrium towards dehybridization of the displacement strand, enabling fluorescence-based target detection through disruption of a fluorophore-quencher pair. The key feature of this design is that it offers two distinct control parameters: displacement strand length (L_DS_) and loop length (L_loop_). This is in contrast to alternative constructs such as aptamer beacons, which have just a single control parameter—displacement strand length—that confers only coarse-grained control over affinity and couples the construct’s kinetics to its thermodynamics^16^. We show mathematically and experimentally that the two control parameters of the ISD design enable us to precisely and independently tune the thermodynamics and kinetics of the resulting aptamer switches. We use this approach to generate an array of aptamer switches that exhibit affinities spanning four orders of magnitude, with equilibrium dissociation constants (*K*_*D*_) ranging from 10 μM to 40 mM and binding kinetics ranging from 170 ms to 3 s—all starting from the same parent ATP aptamer^19,20^. Lastly, we demonstrate that even tighter control of binding affinity and kinetics can be achieved by introducing single-base mismatches into the displacement strand. This approach should be broadly applicable to virtually any aptamer, enabling facile production of highly controllable molecular switches that respond to ligands over a wide range of concentrations and time scales.

## Results

### Design and rationale

The ISD design achieves molecular recognition through concentration-dependent shifts in equilibrium (**Fig. 1**). A fluorophore and a quencher are added to the 5’- and 3’-ends of the construct, respectively, enabling a fluorescent readout of target concentration (**Supplementary Fig. 1a**). Although the thermodynamic and kinetic parameters associated with target binding to the native aptamer 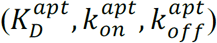 are fixed, we can tune the overall signaling response by altering the parameters of the hybridization/quenching reaction 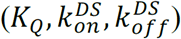.

**Figure 1.**
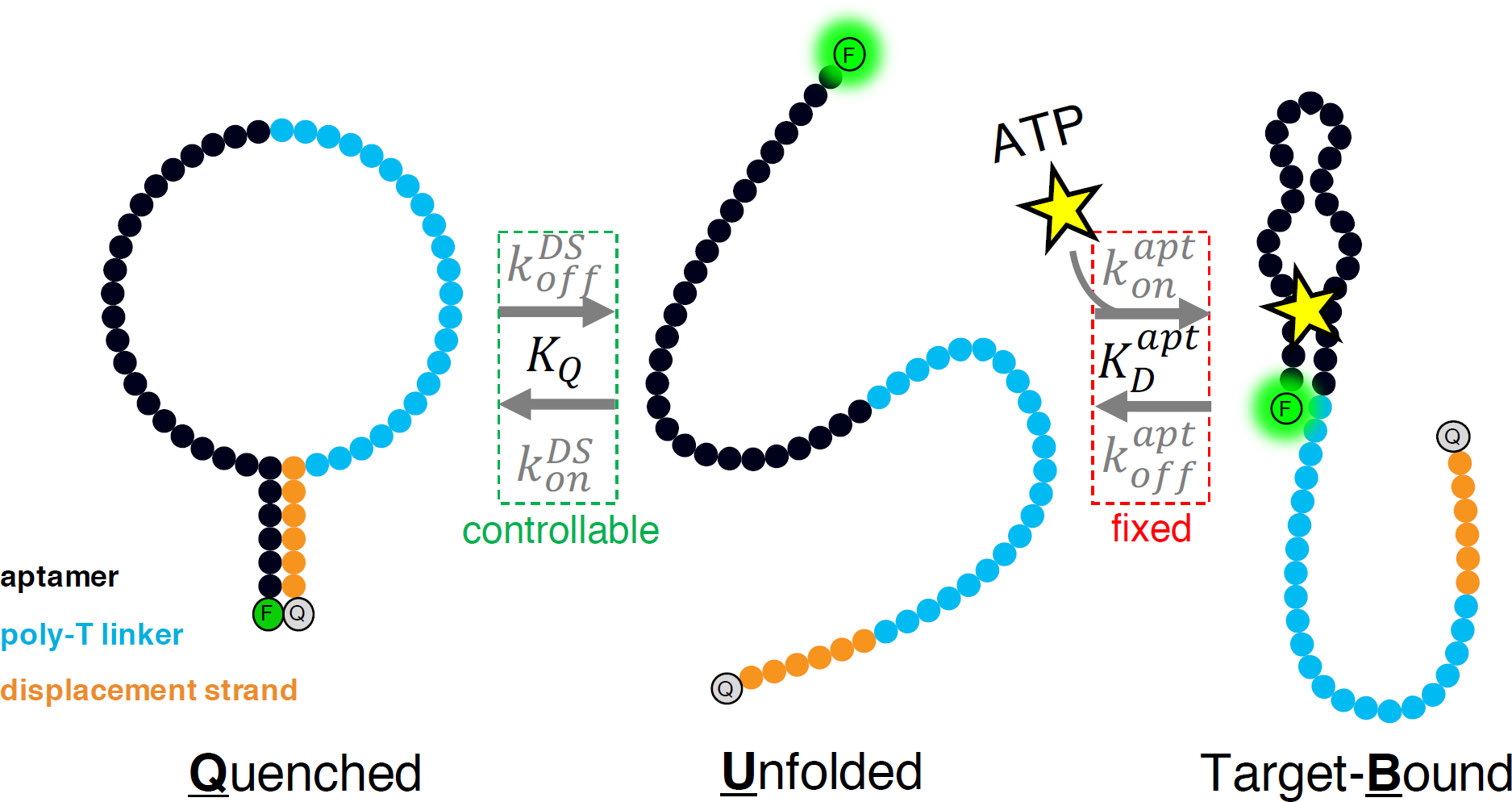
Overview of the intramolecular strand-displacement (ISD) switch design and its relevant parameters. We use an ISD design to convert an existing aptamer into a switch that gives a target-concentration-dependent signal based on the interaction of a fluorophore-quencher pair at the 5’- and 3’- ends of the construct, respectively. In this three-state population shift model, signal is generated by the unfolded and target-bound forms. In the absence of target, the quenched and unfolded states are in equilibrium, defined by K_Q_. Target binding depletes the unfolded population, and the reaction shifts to the right, generating a signal that is proportional to target concentration. We use this model here to generalize the discussion and insights to other aptamers; however, since the ATP aptamer has two binding sites^20^, we have used a modified two-site binding model for fitting and calculations (**Supplementary Fig. 2**, equation S2).

Since hairpin hybridization strength confers coarse-grained control over binding affinity^14^ and linker length confers fine-grained control of binding thermodynamics^15^, the incorporation of both tuning mechanisms makes this switch design highly amenable to the fine-tuning of molecular recognition (**Fig. 2a**). Increasing the hybridization strength of the displacement strand shifts the equilibrium towards the quenched state, which will decrease the background signal at the expense of decreased effective affinity 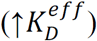 and temporal resolution (**Fig. 2b, c**). Decreasing L_loop_ results in a similar equilibrium shift due to increased effective concentration of the displacement strand, but also results in increased temporal resolution. Independent tuning of the ISD’s thermodynamics and kinetics is made possible by the orthogonal effects of these two parameters.

**Figure 2.**
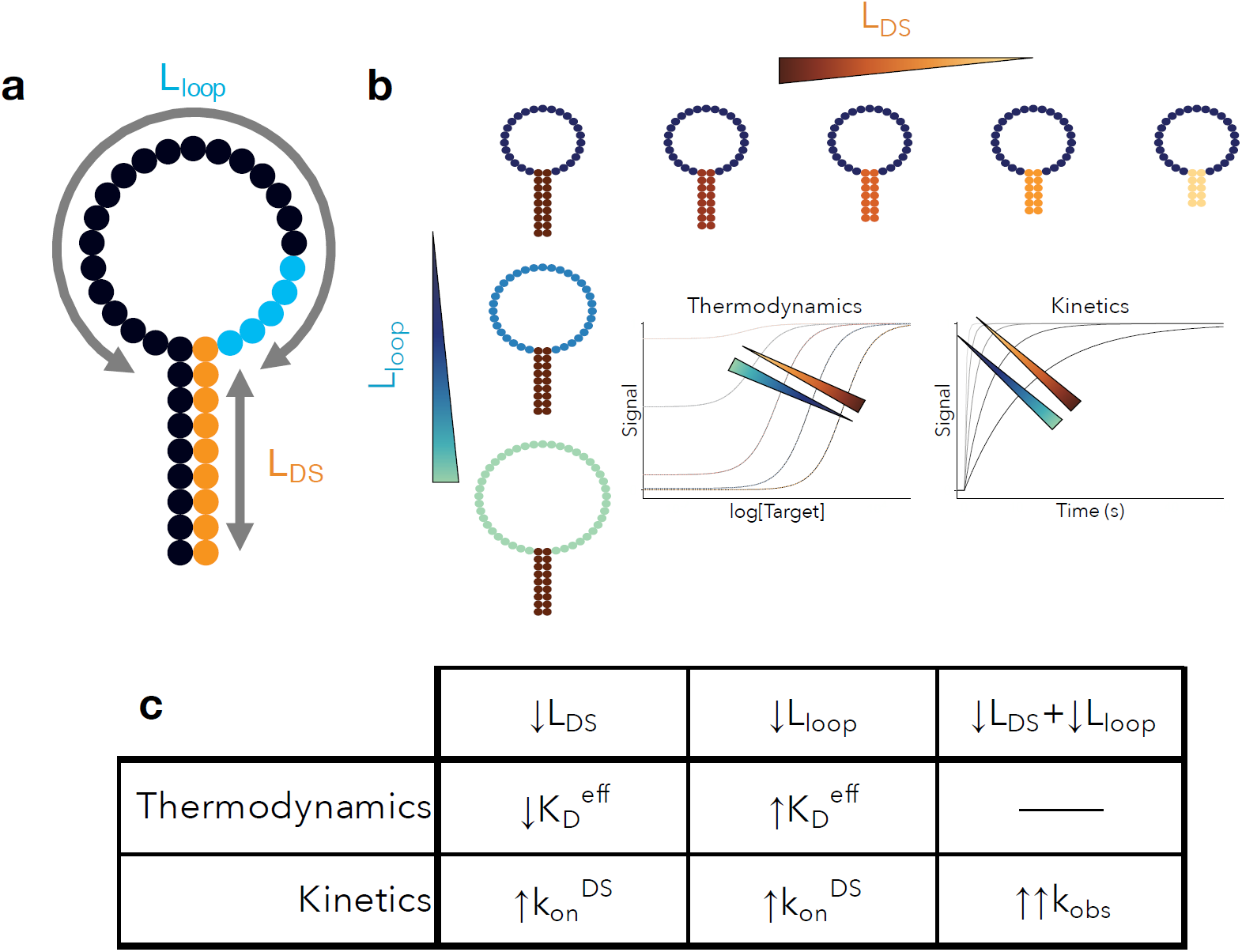
ISD control parameters and their effects. (**a**) By modulating the length of the linker (L_loop_) and the hybridization strength of the hairpin (L_DS_) between the displacement strand and the aptamer, we can control both the kinetics and thermodynamics of our ISD switch. (**b**) On one hand, reducing L_DS_ increases effective binding affinity at the expense of increased background. On the other hand, decreasing L_loop_ decreases background at the expense of lower effective binding affinity. (**c**) Decreasing either parameter increases overall rate of binding, making it possible to increase the kinetics of a given construct while retaining the same K_D_^eff^.

We first used a model system to mathematically test the anticipated effects on binding response, and then confirmed these effects with experimental results from an array of ISD switches (**Supplementary Fig. 3, Supplementary Table 1**) based on a well-studied ATP aptamer^20^ with varying L_loop_ and L_DS_. All of the resulting constructs retain the high selectivity of the native aptamer (**Supplementary Fig. 1b**). We show conclusively that modulating L_loop_ and L_DS_ in tandem decouples the control over the thermodynamics and kinetics of molecular recognition. Lastly, we introduce targeted mismatches as a third tuning parameter to obtain even more precise enthalpic tuning and ultra-fast kinetics over a wide range of binding affinities.

### Principles of ISD molecular switch design

By examining a three-state population shift model, we gain general insights into how the design parameters affect the overall thermodynamics and kinetics of molecular recognition. We have used an induced-fit model for its generalizability and simplicity, but we have also provided derivations for two-site induced-fit and conformational selection (**Supplementary Fig. 2** and **4**, respectively). This inclusion is in recognition of the fact that the ATP aptamer used in this work has two binding sites^20^ and can exhibit both induced-fit and conformational selection behavior^21,22^.

We assume the switch exists in an equilibrium between quenched (*Q*), unfolded (*U*), and target-bound (*B*) states (**Fig. 1**). The distributions of *Q, U*, and *B* are governed by the equilibrium constant (*K*_*Q*_) for the intramolecular quenching reaction,

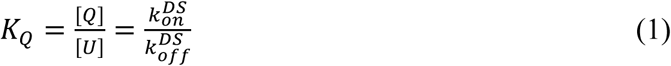

and the dissociation constant of the native aptamer 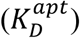,

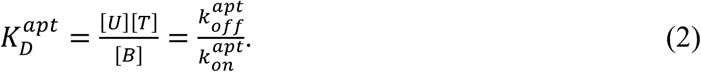

Target binding to the aptamer depletes the unfolded state, shifting the quenching reaction towards the unfolded state, generating more signal. Assuming a quenching efficiency of *η*, this equilibrium shift generates a target concentration-dependent signal given by

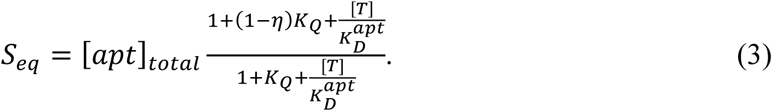

The effective dissociation constant 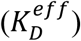, which reflects the effective binding affinity of the overall equilibrium, can be derived^14^ as

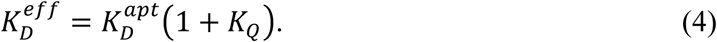

Thus, while the 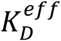 of our construct is defined by the binding properties of the native aptamer 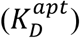, it also depends strongly on *K*_*Q*_. Moreover, the background signal of the construct is also strongly related to *K*_*Q*_ by

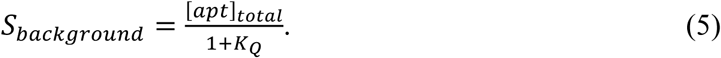

Therefore, it is crucial to understand the relative contributions of L_DS_ and L_loop_ to *K*_*Q*_. We isolate the effects of these independent tuning mechanisms by considering *K*_*Q*_ to be given by

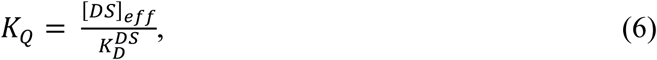

where [*DS*]_*eff*_ is the effective concentration of the displacement strand that arises from covalent coupling to the native aptamer—a function of L_loop_ —and 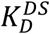 represents the dissociation constant for the hybridization of an unlinked displacement strand—a function of L_DS_. [*DS*]_*eff*_ constitutes the entropic component of *K*_*Q*_, whereas 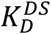 constitutes the enthalpic component of *K*_*Q*_. If the displacement strand is too short (high 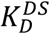) or the linker is too long (low [*DS*]_*eff*_), *K*_*Q*_ will be small, resulting in a large background (Eq. 5) and little signal change upon the addition of target (Eq. 3).

We assume that 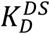 is related to the binding energy of the free, untethered displacement strand, Δ*G*_*DS*_:

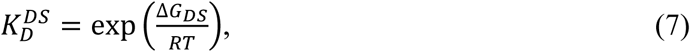

where 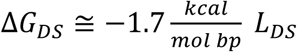 (Ref. 23). We can also make arguments for the approximate scaling of *K*_*Q*_ with L_loop_ based on observations of rates of DNA hairpin closure as a function of loop size. Since the dissociation rate 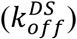 is relatively independent of L_loop_ and the association rate 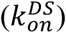 has been shown to scale inversely with L_loop_ to the power of 2.6 ± 0.3 (Ref. 24), we can approximate that 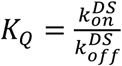 scales as ∼L_loop_^−2.6^. Therefore, increases in L_DS_ or decreases in L_loop_ will be mirrored by an increase in *K*_*Q*_, shifting the equilibrium towards the quenched state, which results in decreased background signal at the expense of higher 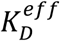. Moreover, we can derive that L_loop_ will have a subtler per-base effect on *K*_*Q*_ relative to L_DS_ and that increases in L_loop_ will have diminishing returns because 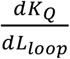 decreases as 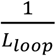 (**Supplementary Calculation 1**).

This qualitative reasoning also yields testable insights into the kinetic control of the system. We have discussed how increases in L_loop_ decrease 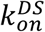 with negligible effects on 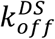, and others have shown that increasing hybridization strength decreases 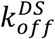 much more than 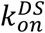 (Ref. 26). Since the observed binding rate (*k*_*obs*_) depends on the sum of 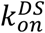 and 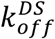 as

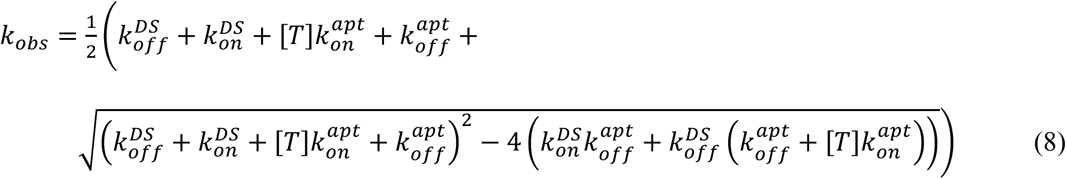

(derived in SI, **Supplementary Calculation 2**), we can determine that decreasing L_loop_ or L_DS_ will both increase *k*_*obs*_. To summarize, we expect that L_DS_ and L_loop_ will have opposing effects on ISD switch thermodynamics but additive effects on the kinetics, and that L_DS_ will have a more profound impact per base than L_loop_ on effective binding affinity.

### Experimental characterization of the ISD switch

In order to experimentally validate the predictions of this model, we characterized ligand binding for an array of ISD switches (**Supplementary Fig. 3**) derived from the same ATP aptamer^20^. We introduced displacement strands with L_DS_ ranging from 5–9 base pairs (bp) and poly-T linkers of various lengths to yield L_loop_ ranging from 23–43 nucleotides (nt). As expected, increasing L_DS_ with a constant L_loop_ resulted in decreased background signal and lower apparent affinity 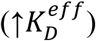 (**Fig. 3a**). Fits of equation S1 (**Supplementary Fig. 2**) reveal a clear trend in which 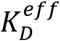 increases with L_DS_, reflecting a reduction in effective binding affinity (**Fig. 3b**). We observed that 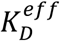 can be decreased up to 1,200-fold by removing three bases from the displacement strand (*e.g.*, converting 9_23 to 6_23), with an average fold change of ∼6.7 ± 2.4 per base. Notably, the addition or removal of a single base from the displacement strand can shift 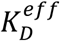 by more than an order of magnitude.

**Figure 3.**
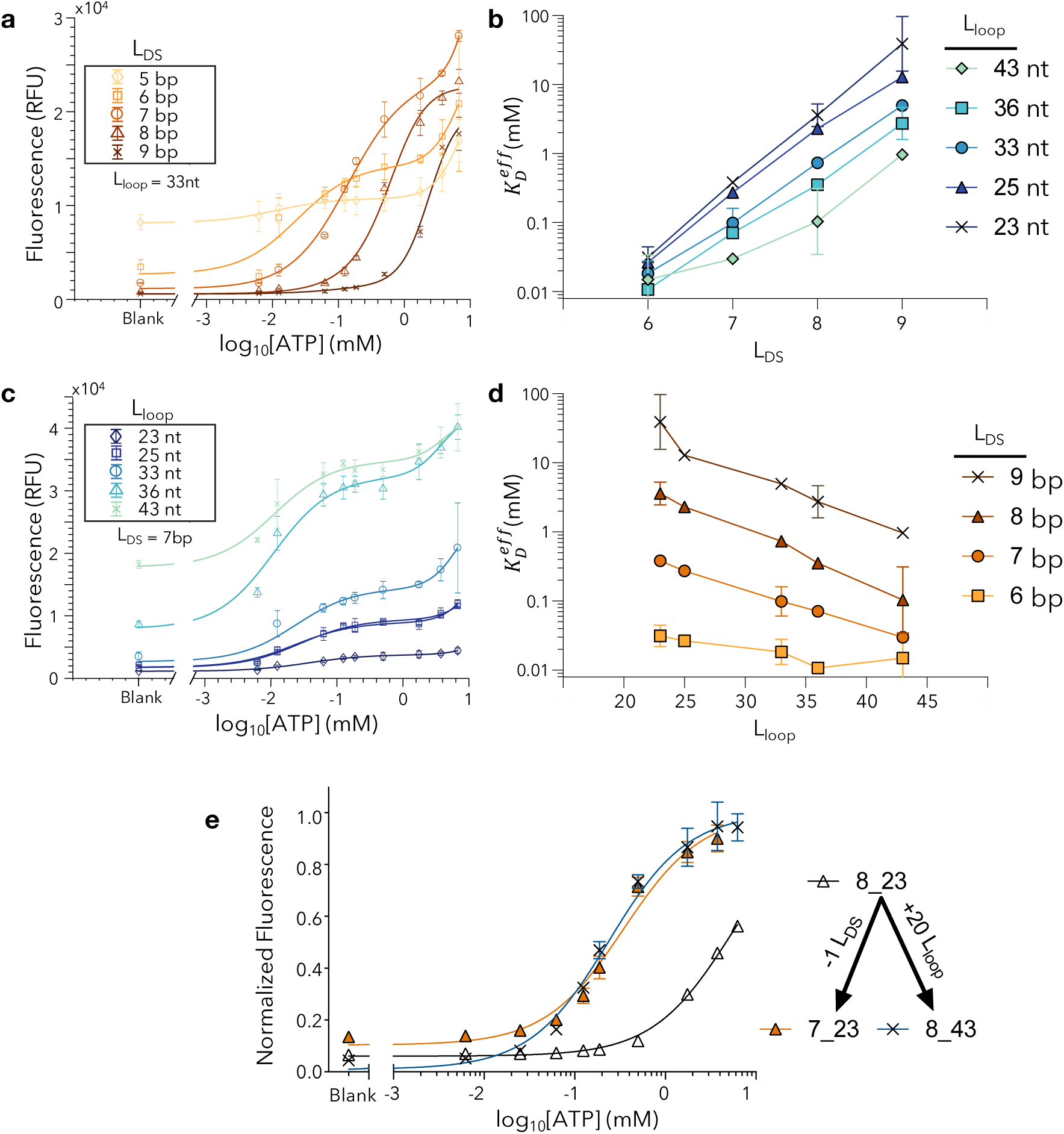
Binding curve modulation via ISD switch design. (**a**) Changing L_DS_ from 5 to 9 nt while maintaining L_loop_ at 33 nt shifts the binding curve to the right and reduces background signal. (**b**) 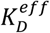 increases with L_DS_ given a fixed L_loop_. (**c**) Changing L_loop_ from 23 to 43 nt while holding L_DS_ constant at 7 bp shifts the binding curve to the left and increases background signal. (**d**) 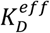 decreases with L_loop_ given a fixed L_DS_. (**e**) The removal of a single base from the displacement strand of 8_23, generating 7_23, causes the same binding curve shift as adding 20 bases to the linker of 8_23, generating 8_43. All plots are averaged over three replicates. Error bars in **a, c**, and **e** represent the standard deviation of the average; error bars in **b** and **d** represent the error calculated via propagation of errors (Methods). Fits in **a** and **c** were conducted according to equation S1 (**Supplementary Fig. 2**). **a** and **c** show raw fluorescence data, whereas panel **e** is normalized by a single-site hyperbolic binding curve. Fit parameters for constructs in which L_DS_ = 5 have been omitted from **b** and **d** because we were unable to obtain robust fits for the parameters. Raw thermodynamic plots for all ISD constructs is provided in **Supplementary Fig. 5**.

In contrast, we observed that changing L_loop_ has a subtler per-base effect on 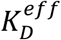, with just a ∼0.83 ± 0.15 fold decrease in 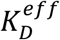 per additional base (**Fig. 3c, d**). On average, the linker must be increased by 17.7 ± 11.9 nt in order to shift the binding curve by an amount equivalent to the removal of a single base to the displacement strand. This loop/base equivalence value varies from construct to construct (**Supplementary Calculation 1**) but is epitomized by the observation that adding 20 nt to the linker of 8_23 (generating 8_43) results in the same 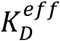 as removing 1 bp from the displacement strand of 8_23 (generating 7_23) (**Fig. 3e**). The vast difference in these effects enables us to modulate effective binding affinity both finely (by tuning L_loop_) and over a wide functional range (by tuning L_DS_).

Next, we measured the temporal response of molecular recognition for all of our constructs to validate the previously described kinetic contributions of L_loop_ and L_DS_. As anticipated, we found that decreasing L_DS_ with a constant L_loop_ (**Fig. 4a, b**) or decreasing L_loop_ with a constant L_DS_ (**Fig. 4c, d**) resulted in faster temporal responses. By combining these two tuning mechanisms, we could vary the switching time constant (*k*_*obs*_^−1^) by over 20-fold, ranging from ∼3 s to ∼170 ms. We note that even our slowest constructs represent a marked improvement over traditional aptamer beacons, which typically exhibit time constants on the order of minutes to hours^16,27^. The fast kinetics of the ISD switch are attributable to a much higher 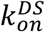 resulting from the high effective concentration of the displacement strand. This high 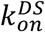 allows us to use much shorter displacement strands than are possible with aptamer beacons, which in turn results in a much faster 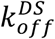. The simultaneous increase in both 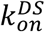 and 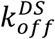 greatly increases *k*_*obs*_ (Eq. 8). Indeed, switches with L_DS_ = 5 achieved temporal responses exceeding the time resolution of our detector; with a time delay of 465 ms between injection and measurement, 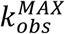 is approximately 10 *s*^−1^.

**Figure 4.**
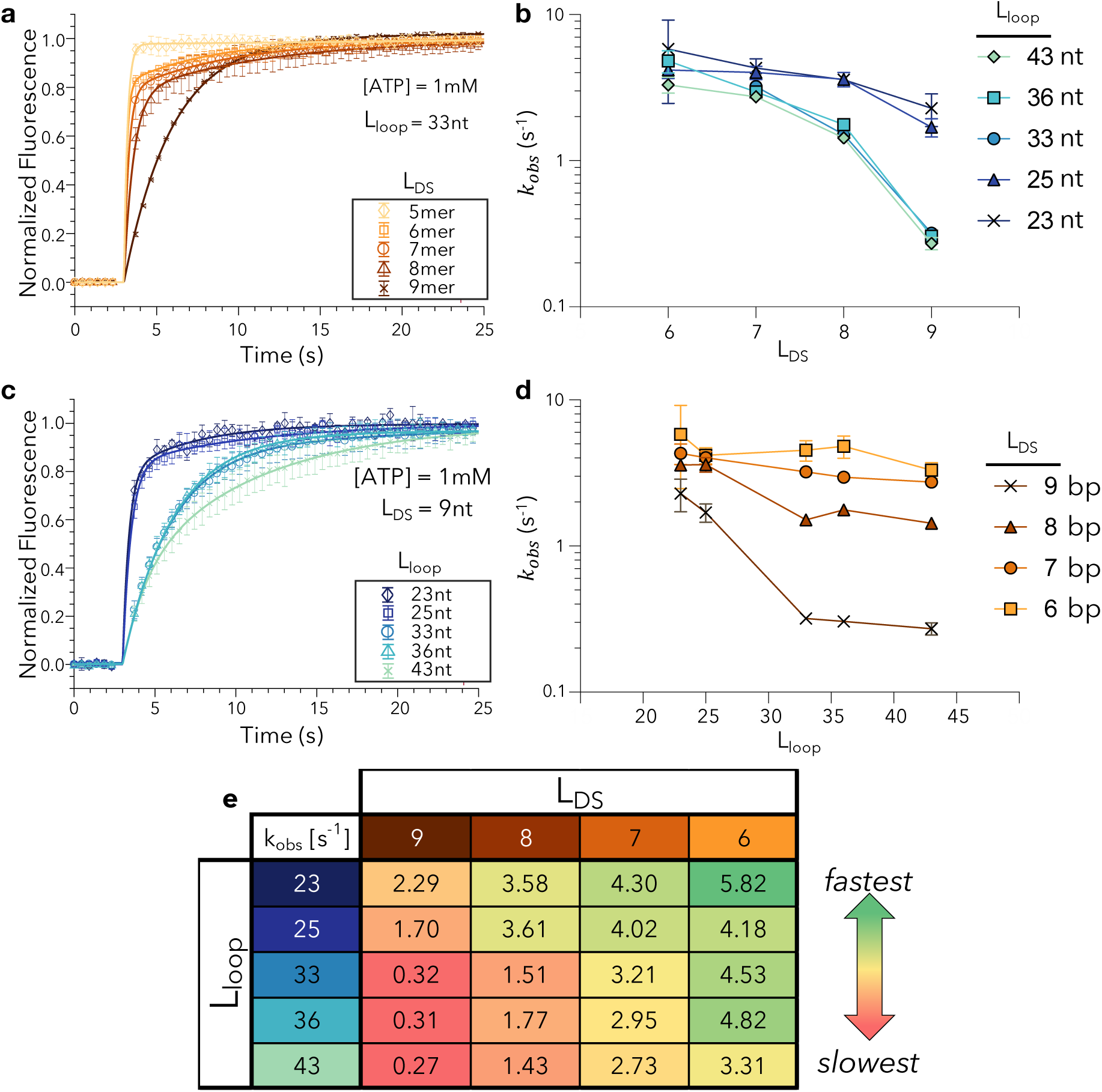
Temporal response modulation via ISD switch design. (**a**) Normalized signal change upon injection of 1 mM ATP at 3.5 s. Increasing L_loop_ while holding L_DS_ constant at 33 nt results in slower kinetics. (**b**) Effect on *k*_*obs*_ of increasing L_loop_ in an ISD construct with constant L_DS_ in the presence of 1 mM ATP. (**c**) Normalized signal change upon injection with 1 mM ATP at 3.5 s. Increasing L_loop_ while keeping L_DS_ constant at 9 results in slower kinetics. (**d**) Effect on *k*_*obs*_ of increasing L_DS_ in an ISD construct with constant L_loop_ in the presence of 1 mM ATP. (**e**) *k*_*obs*_ as a function of both L_DS_ and L_loop_ after injection with 1 mM ATP. Sequences with L_DS_ = 5 had switching responses faster than the time resolution of our detector and could not be fit accurately and have thus been omitted from panels **b, d**, and **e**. Error bars in **a** and **c** represent standard deviation over three replicates, whereas those in **b** and **d** represent the 95% confidence intervals in the fit parameter. Confidence intervals for **e** are listed in **Supplementary Fig. 3**.

### Decoupling thermodynamics and kinetics

Our thermodynamic and kinetic findings introduce the possibility of designing ISD switches such that temporal resolution can be tuned completely independently of binding affinity. Since the effective affinity of our construct depends on the equilibrium constant for hairpin formation, 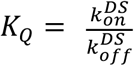, it is clear that the binding kinetics can be increased while maintaining the same effective affinity provided the ratio between association and dissociation rates is preserved. Decreasing L_loop_ increases the binding rate and decreases the effective affinity of our construct. This decrease in affinity can be offset by shortening L_DS_, which in turn results in a net additive increase in temporal resolution. Based on the observed dependencies of our two control parameters, we hypothesized that it should be feasible to achieve faster switching responses and maintain effective affinity by decreasing L_loop_ and L_DS_ simultaneously.

We confirmed this prediction with three pairs of constructs (**Fig. 5a**) that each have statistically indistinguishable effective affinities (**Fig. 5b**). Constructs with both a longer loop length and a stronger hairpin had universally slower temporal resolution (**Fig. 5c**). Tandem tuning of the two parameters allowed us to increase the aptamer temporal response by up to 6-fold without changing 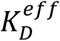. For example, in pair I, decreasing L_loop_ from 36- to 23-nt increases 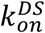 while decreasing L_DS_ from 9- to 8-bp increases 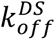, resulting in much more rapid observed kinetics with a roughly constant *K*_*Q*_. Importantly, we were able to achieve this tunability over a wide range of effective affinities (∼90 µM to ∼5.8 mM).

**Figure 5.**
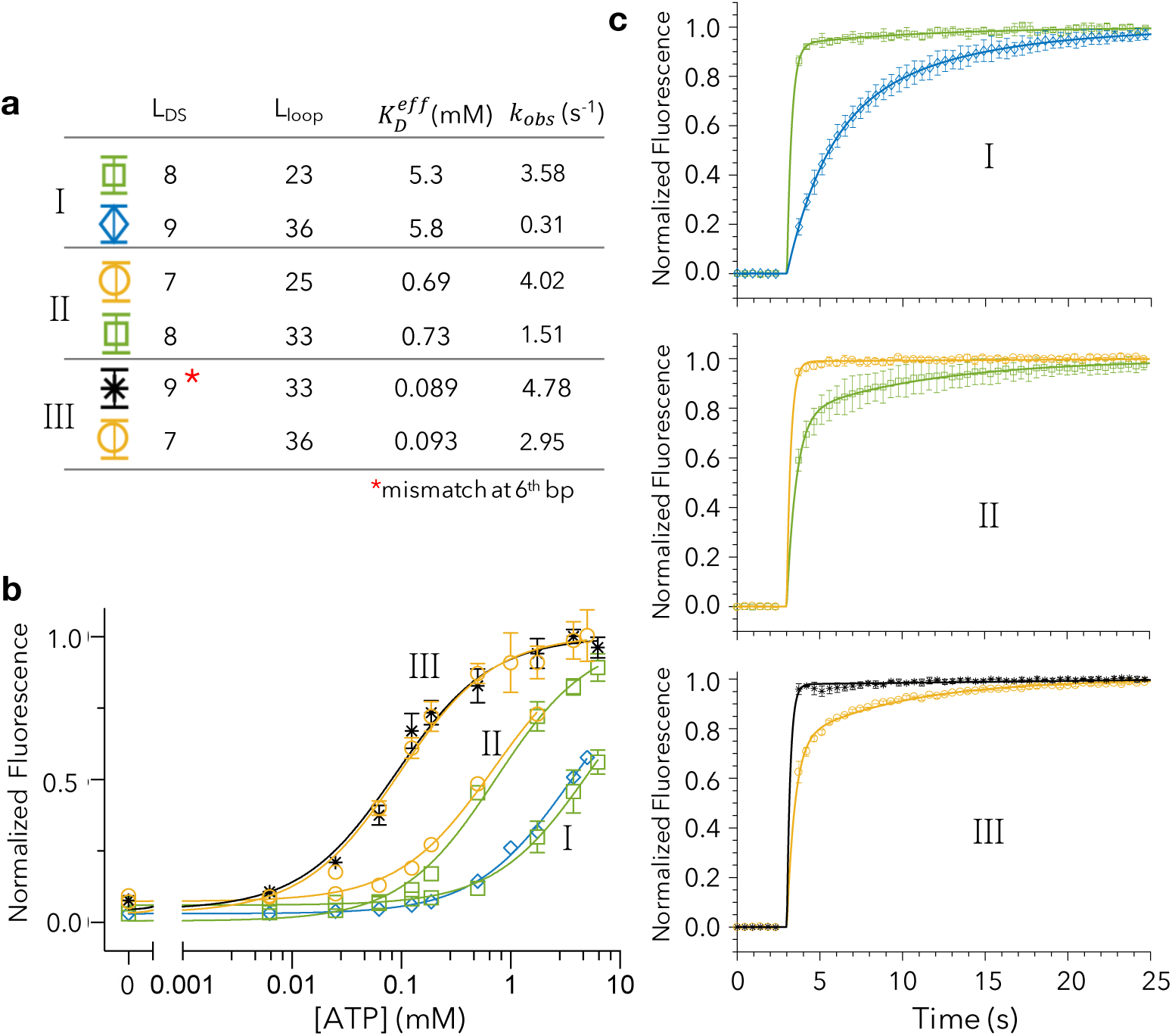
Independent tuning of kinetics and thermodynamics through simultaneous changes to L_DS_ and L_loop_. Increasing both L_DS_ and L_loop_ simultaneously has synergistic effects on temporal resolution, but opposing effects on 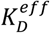. Therefore it should be possible to change the kinetic response of the switches while holding the affinity constant **(a)** We confirmed this by examining three pairs (I, II, III) of ISD constructs. **(b)** Each pair exhibited nearly identical binding curves, **(c)** but has been engineered in terms of L_DS_ and L_loop_ to exhibit vastly different kinetic responses. The 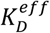 listed in **a** and the binding curve fits in **b** are respectively derived from and normalized to a single-site hyperbolic binding curve. For kinetics measurements, pair I was run at [ATP] = 2.5 mM and pairs II and III were run at [ATP] = 1 mM. Plots were averaged over three replicates.

### Precision tuning through mismatches in the displacement strand

We hypothesized that even finer control over the hybridization strength of the duplexed region of the ISD construct should be possible if we manipulate the complementarity of the two strands. Indeed, by expanding our design space to include single-base mismatches, we calculated that we could increase the theoretical resolution of our tuning capability by more than 10-fold relative to that of perfectly-matched displacement strands (**Supplementary Fig. 6a**). Since the introduction of mismatches has been shown to drastically increase k_off_ and k_on_ for two hybridizing strands^26^, we anticipated that the introduction of mismatches would greatly increase the observed kinetics. Thus, mismatches should enable finer enthalpic control over the binding curve and enhance our ability to increase kinetics independently of affinity.

To experimentally confirm these predictions, we introduced single mismatches of different identities (A, G, C or T) at various positions throughout the displacement strands of three constructs: 8_33 (L_DS_ = 8, L_loop_ = 33), 9_33 (L_DS_ = 9, L_loop_ = 33), and 10_33 (L_DS_ = 10, L_loop_ = 33). Upon comparing the thermodynamic properties of the original constructs to those containing the mismatches, we found that we were able to obtain more closely spaced binding curves based on the position and identity of the mismatch. For constructs with a 33-nt loop, for example, the average fold change between the set of 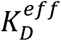 values that are obtainable from perfectly matched displacement strands was 6.55 ± 1.01 (**Fig. 6a**). However, using just a small subset of all the possible mismatches, we were able to reduce this average spacing to 1.72 ± 0.34 (**Fig. 6b; Supplementary Fig. 6b**). Therefore, by modulating the position and identity of mismatches, we can generate sets of constructs that yield much finer enthalpic control than would be possible by changing L_DS_ alone. Lastly, the incorporation of mismatches substantially increases 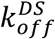 (Ref. 26), such that mismatches not only confer greater control over the thermodynamics but also dramatically increase temporal resolution relative to perfectly-matched displacement strands (**Supplementary Fig. 6c**).

**Figure 6.**
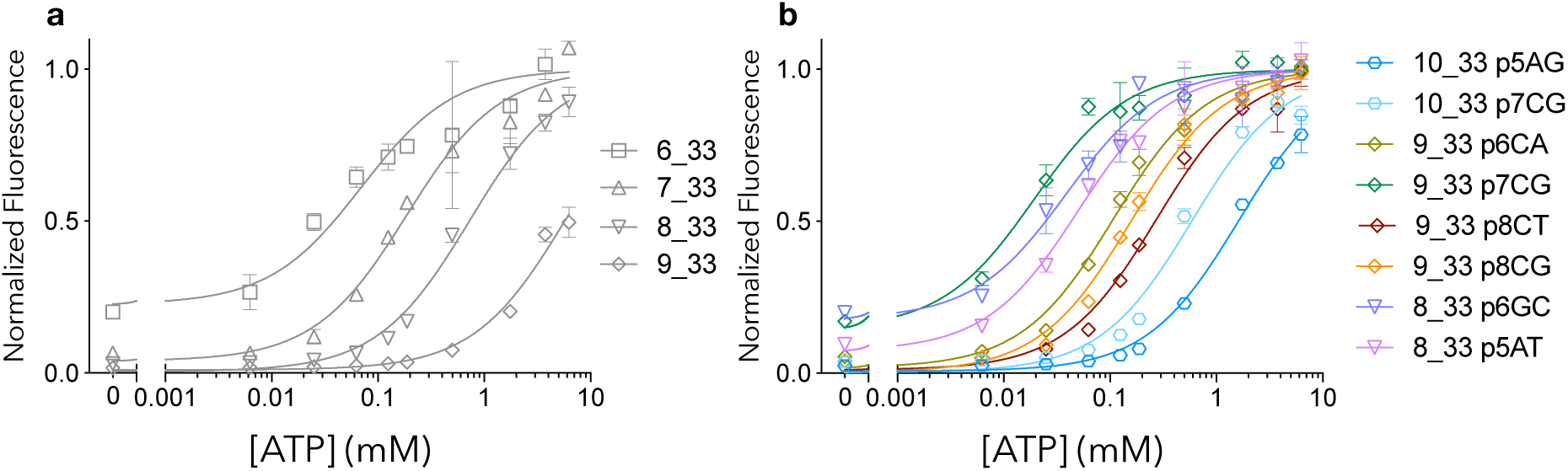
Fine tuning of binding curves via the incorporation of mismatches in the displacement strand. (**a**) Modulating affinity via L_DS_ alone leads to huge jumps in affinity. **(b)** In contrast, the incorporation of single mismatches into the displacement strand produces much more granular shifts in effective binding affinity. Mismatches are described such that “9_33 p6CG” indicates that position 6, as defined from the 3’- end, was changed from C to G. Curves were averaged over three replicates and were fit to and normalized by a single-site hyperbolic binding curve.

## Conclusion

Although aptamer-based molecular switches are powerful tools in biotechnology, their utility has been constrained by a limited ability to rationally engineer their binding characteristics in terms of affinity and kinetics. Prior studies have indicated that the thermodynamics and kinetics of such switches are coupled in such a manner that gains in one parameter will result in sacrifices in the other^16^. Here, we have demonstrated an aptamer switch design that allows remarkably precise independent control of its thermodynamic and kinetic parameters. Our ISD construct connects an existing aptamer to a partially complementary displacement strand via a poly-T linker, such that alterations in the length of either feature can meaningfully shift the equilibrium binding to the aptamer’s target. We used a mathematical model to demonstrate how changes in the hybridization strength of the displacement strand would confer coarse control over switch affinity; at the same time, changes in linker length produce a subtler effect per added or removed base. We subsequently confirmed this expectation experimentally and demonstrated the capacity to carefully manipulate the binding characteristics of our switch through these two parameters. For example, we can increase binding kinetics by an order of magnitude with minimal effect on aptamer affinity by selectively shortening both L_DS_ and L_loop_. Furthermore, we have shown that even finer tuning of ISD binding properties is possible when we manipulate the strength of displacement strand hybridization through the targeted introduction of individual base-mismatches into the displacement strand sequence.

As the desire for rapid molecular detection becomes more prevalent, so too will the need to tune the kinetics of molecular recognition independently of binding affinity. Our approach is advantageous in this regard, as it offers opportunities for control that exceed those of existing molecular switch designs, which are generally constrained by tight coupling of kinetic and thermodynamic parameters and offer less freedom for structural manipulation. We have demonstrated the feasibility of achieving ultrafast kinetic responses (on the order of hundreds of milliseconds) with our ISD constructs without meaningfully sacrificing target affinity, whereas aptamer beacons typically exhibit kinetics on the order of minutes to hours. Critically, our molecular switch design should be compatible with virtually any aptamer sequence, making it feasible to design optimized molecular switches that are ideally suited for a diverse array of biotechnology and synthetic biology applications.

## Methods

### Reagents

All chemicals were purchased from Thermo Fisher Scientific unless otherwise noted, including ATP (25 μmol, 100 mM), Tris-HCl Buffer (1 M, pH 7.5), magnesium chloride (1 M, 0.2 μm filtered), and Hyclone molecular biology-grade water (nuclease-free). Oligonucleotides modified with Cy3 fluorophore at the 5’ ends and DABCYL quencher at the 3’ ends, purified by HPLC, were purchased from Integrated DNA Technologies. All sequences used in this work are shown in **Supplementary Table 1**. All oligonucleotides were resuspended in nuclease-free water and stored at −20 °C.

### Measurements of Effective Binding Affinity

To obtain binding curves, 40 µL reactions were prepared in 1x ATP binding buffer (10 mM Tris-HCl, pH 7.5 and 6 mM MgCl_2_) with 250 nM aptamer and final ATP concentrations in the range of 6.25 μM to 6.75 mM. The fluorescence spectra for all samples were measured at 25 °C on a Synergy H1 microplate reader (BioTeK). Emission spectra were monitored in the 550–700 nm range with Cy3 excitation at 530 nm and a gain of 100, in 96-well plates. All measurements were performed in triplicate. Representative concentration-dependent emission spectra are shown in **Supplementary Figure 1**.

### Measurements of Binding Kinetics

ISD constructs of varying linker and displacement strand lengths were suspended at a concentration of 333.3 nM in a 30 μL total volume of 1x ATP binding buffer (10 mM Tris-HCl, pH 7.5 and 6 mM MgCl_2_). Kinetic fluorescence measurements of the quencher-fluorophore pair were made using a Synergy H1 microplate reader. Cy3 was excited at 530 nm, and unquenched fluorescence was measured at 570 nm emission using monochromators at the minimum possible regular time interval of 0.465 seconds. After timed injection of 10 µL ATP (final [ATP] = 0, 1, or 2.5 mM) in 1x ATP binding buffer into the 30 μL ISD solution, we measured the kinetic response in the 40 μL sample volume. We first normalized all kinetic data relative to the 0 μM target concentration to account for the effect of sample volume change upon injection with ATP. For plotting, we normalized the curves to range from 0 to 1 in order to visually emphasize changes in rate constants rather than the background and peak levels that are dictated by the thermodynamics. All measurements were performed in triplicate.

### Thermodynamic Analysis

Three replicates of inputs (*X* = log([*ATP*]) in mM) versus outputs (*Y* in raw RFU intensity) were fit individually for each construct to extract the effective binding affinity. The resultant parameters from fitting *X* and *Y* to equation S2 were averaged over the three independent fits. We fit the logarithmic values of thermodynamic constants *pK*_*D*1_ = − log(*K* _*D*1_), *pK*_*D*2_ = − log(*K*_*D*2_), and *pK*_*Q*_ = − log(*K*_*Q*_) such that equation S2 becomes:

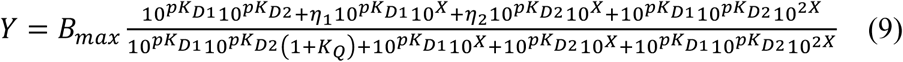

Fits were performed via MATLAB’s *lsqcurve* function with **initial guesses** equal to

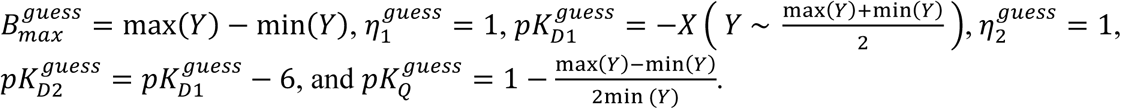

**Upper bounds** were set to 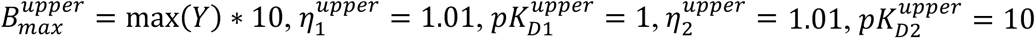, and 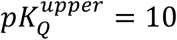.

**Lower bounds** were set to 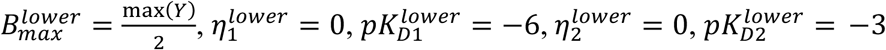, and 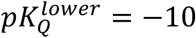.

Fitting was performed using a maximum of 100,000 iterations. *pK*_*D*1_ and *pK*_*Q*_ were averaged over at least three replicate fits for each construct. The effective binding affinity, 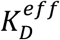, was then calculated by

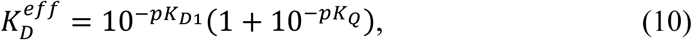

with the standard error given by propagation of errors

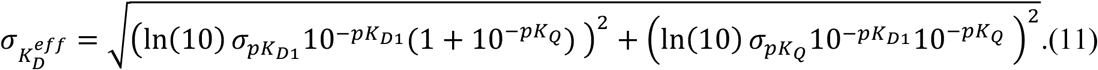

The curves plotted in **Figure 3a** and **c** were fit to the average of the three replicates.

### Normalized Thermodynamic Plots

For ease of comparison, the data in **Fig. 3e, Fig. 5b**, and **Fig. 6** were fit to and normalized according to a single-site hyperbolic binding curve:

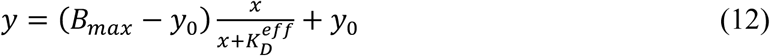

For **Fig. 3e, Fig. 5b**, and **Fig. 6**, the average fluorescence value was divided by B_max_ prior to plotting. Error bars were reported via propagation of errors for the standard deviation of the average fluorescence and the error in the fit for B_max_.

### Kinetic Analysis

Kinetic data were first normalized to a zero ATP control to account for changes in volume due to the injection of ATP. We fit each replicate individually to

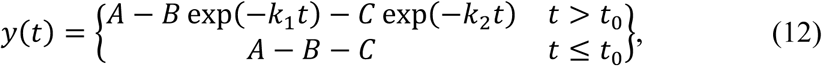

where k_1_>k_2_.

Each replicate was then normalized to a range of zero to one by:

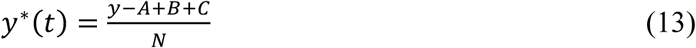

where 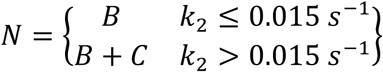. A piecewise function was used for N to control for an artifact of the fitting function, wherein if there is no observable *k*_*2*_, the fit function still forces the fit to *k*_*2*_ which can result in extremely large values of C. Therefore, we omit C from the normalization if *k*_*2*_ is very slow. The normalized responses were averaged and the average response was again fit to equation 12 to obtain a rate representative of all three trials. Rate constants are reported as the best fit values ± 95% upper/lower confidence intervals.

## Data availability

All data are available from the authors upon reasonable request.

## Acknowledgements

This work was supported by NIH SPARC Initiative (OT2OD025342), NIH (1R01GM129313), Wu Tsai Neurosciences Institute at Stanford, Stanford Maternal and Child Health Research Institute and the Chan-Zuckerberg Biohub. H.T.S is a Chan Zuckerberg Biohub investigator. We are thankful to Philips Healthcare and NSERC PDF (A.A.H.), and the Medtronic Foundation Stanford Graduate Fellowship (I.A.P.T) We also thank Evelin Sullivan of the Technical Communications Program at Stanford for her thoughtful comments and edits on the manuscript.

## Author contributions

B.D.W., A.A.H., and H.T.S. conceived the initial concept. B.D.W., A.A.H., and I.A.P.T. designed experiments. A.A.H. and I.A.P.T. executed experiments. B.D.W. developed the model and analyzed the data. B.D.W., M.E., and H.T.S. wrote the manuscript. All authors edited, discussed, and approved the whole paper.

## Additional information

### Online Supplementary Information

accompanies this paper

