## Supplemental Information for "Independent Control of the Thermodynamic and Kinetic Properties of Aptamer Switches"

### Contents

**Supplementary Table 1 l** DNA sequences used in this work
**Supplementary Figure 1 l** Raw fluorescence traces and specificity test
**Supplementary Figure 2 l** Two-site induced fit model
**Supplementary Figure 3 l** Summary of thermodynamic and kinetic results
**Supplementary Figure 4 l** Conformational selection model
**Supplementary Figure 5 l** Raw binding curves for all constructs
**Supplementary Figure 6 l** Additional mismatch results
**Supplementary Calculation 1 l** Relative effects of linker length and displacement strand length on binding affinity
**Supplementary Calculation 2 l** Derivation of signaling kinetics for induced fit

**Supplementary Table 1 | Sequences used in this work.** Black represents the aptamer sequence, blue represents the poly-T linker, and orange represents the displacement strand. Underlined sequence is complementary to the displacement strand. $L_{loop}$ is equal to the length of the linker plus the number of bases between the underlined region and the displacement strand. The loop length in this design cannot be shorter than that of the aptamer minus the length of the displacement strand (in this case, 23 nt). Mismatches introduced into the ISD constructs are shown in red.

| | $L_{DS}$ | $L_{loop}$ | Sequence 5' --> 3' | Total Length |
|---|---|---|---|---|
| Perfect match | 10 | 33 | CACCTGGGGGAGTATTGCGGAGGAAGGTTTTTTTTTTTTTTTTCCCCCAGGTG | 53 |
| | 9 | 23 | CACCTGGGGGAGTATTGCGGAGGAAGGTTTTTCCCCAGGTG | 41 |
| | 9 | 25 | CACCTGGGGGAGTATTGCGGAGGAAGGTTTTTTTCCCCAGGTG | 43 |
| | 9 | 33 | CACCTGGGGGAGTATTGCGGAGGAAGGTTTTTTTTTTTTTTTCCCCAGGTG | 51 |
| | 9 | 36 | CACCTGGGGGAGTATTGCGGAGGAAGGTTTTTTTTTTTTTTTTTTCCCCAGGTG | 54 |
| | 9 | 43 | CACCTGGGGGAGTATTGCGGAGGAAGGTTTTTTTTTTTTTTTTTTTTTTTTTCCCCAGGTG | 61 |
| | 8 | 23 | CACCTGGGGGAGTATTGCGGAGGAAGGTTTTCCCAGGTG | 39 |
| | 8 | 25 | CACCTGGGGGAGTATTGCGGAGGAAGGTTTTTTCCCAGGTG | 41 |
| | 8 | 33 | CACCTGGGGGAGTATTGCGGAGGAAGGTTTTTTTTTTTTTTCCCAGGTG | 49 |
| | 8 | 36 | CACCTGGGGGAGTATTGCGGAGGAAGGTTTTTTTTTTTTTTTTTCCCAGGTG | 52 |
| | 8 | 43 | CACCTGGGGGAGTATTGCGGAGGAAGGTTTTTTTTTTTTTTTTTTTTTTTTCCCAGGTG | 59 |
| | 7 | 23 | CACCTGGGGGAGTATTGCGGAGGAAGGTTTCCAGGTG | 37 |
| | 7 | 25 | CACCTGGGGGAGTATTGCGGAGGAAGGTTTTTCCAGGTG | 39 |
| | 7 | 33 | CACCTGGGGGAGTATTGCGGAGGAAGGTTTTTTTTTTTTTCCAGGTG | 47 |
| | 7 | 36 | CACCTGGGGGAGTATTGCGGAGGAAGGTTTTTTTTTTTTTTTTCCAGGTG | 50 |
| | 7 | 43 | CACCTGGGGGAGTATTGCGGAGGAAGGTTTTTTTTTTTTTTTTTTTTTTTCCAGGTG | 57 |
| | 6 | 23 | CACCTGGGGGAGTATTGCGGAGGAAGGTTCAGGTG | 35 |
| | 6 | 25 | CACCTGGGGGAGTATTGCGGAGGAAGGTTTTCAGGTG | 37 |
| | 6 | 33 | CACCTGGGGGAGTATTGCGGAGGAAGGTTTTTTTTTTTTCAGGTG | 45 |
| | 6 | 36 | CACCTGGGGGAGTATTGCGGAGGAAGGTTTTTTTTTTTTTTTCAGGTG | 48 |
| | 6 | 43 | CACCTGGGGGAGTATTGCGGAGGAAGGTTTTTTTTTTTTTTTTTTTTTTCAGGTG | 55 |
| | 5 | 23 | CACCTGGGGGAGTATTGCGGAGGAAGGTAGGTG | 33 |
| | 5 | 25 | CACCTGGGGGAGTATTGCGGAGGAAGGTTTAGGTG | 35 |
| | 5 | 33 | CACCTGGGGGAGTATTGCGGAGGAAGGTTTTTTTTTTTAGGTG | 43 |
| | 5 | 36 | CACCTGGGGGAGTATTGCGGAGGAAGGTTTTTTTTTTTTTTAGGTG | 46 |
| | 5 | 43 | CACCTGGGGGAGTATTGCGGAGGAAGGTTTTTTTTTTTTTTTTTTTTTAGGTG | 53 |
| Mismatches | 10 | 33 | CACCTGGGGGAGTATTGCGGAGGAAGGTTTTTTTTTTTTTTTTCGCCCAGGTG | 53 |
| | 10 | 33 | CACCTGGGGGAGTATTGCGGAGGAAGGTTTTTTTTTTTTTTTTCCCGCAGGTG | 53 |
| | 10 | 33 | CACCTGGGGGAGTATTGCGGAGGAAGGTTTTTTTTTTTTTTTTCCCCCTGGTG | 53 |
| | 10 | 33 | CACCTGGGGGAGTATTGCGGAGGAAGGTTTTTTTTTTTTTTTTCCCCCGGGTG | 53 |
| | 9 | 33 | CACCTGGGGGAGTATTGCGGAGGAAGGTTTTTTTTTTTTTTTCGCCAGGTG | 51 |
| | 9 | 33 | CACCTGGGGGAGTATTGCGGAGGAAGGTTTTTTTTTTTTTTTCTCCAGGTG | 51 |
| | 9 | 33 | CACCTGGGGGAGTATTGCGGAGGAAGGTTTTTTTTTTTTTTTCACCAGGTG | 51 |
| | 9 | 33 | CACCTGGGGGAGTATTGCGGAGGAAGGTTTTTTTTTTTTTTTCCGCAGGTG | 51 |
| | 9 | 33 | CACCTGGGGGAGTATTGCGGAGGAAGGTTTTTTTTTTTTTTTCCCGAGGTG | 51 |
| | 9 | 33 | CACCTGGGGGAGTATTGCGGAGGAAGGTTTTTTTTTTTTTTTCCCTAGGTG | 51 |
| | 9 | 33 | CACCTGGGGGAGTATTGCGGAGGAAGGTTTTTTTTTTTTTTTCCCAAGGTG | 51 |
| | 8 | 33 | CACCTGGGGGAGTATTGCGGAGGAAGGTTTTTTTTTTTTTTCGCAGGTG | 49 |
| | 8 | 33 | CACCTGGGGGAGTATTGCGGAGGAAGGTTTTTTTTTTTTTTCCGAGGTG | 49 |
| | 8 | 33 | CACCTGGGGGAGTATTGCGGAGGAAGGTTTTTTTTTTTTTTCCCTGGTG | 49 |
| | 8 | 33 | CACCTGGGGGAGTATTGCGGAGGAAGGTTTTTTTTTTTTTTCCCACGTG | 49 |

**a**

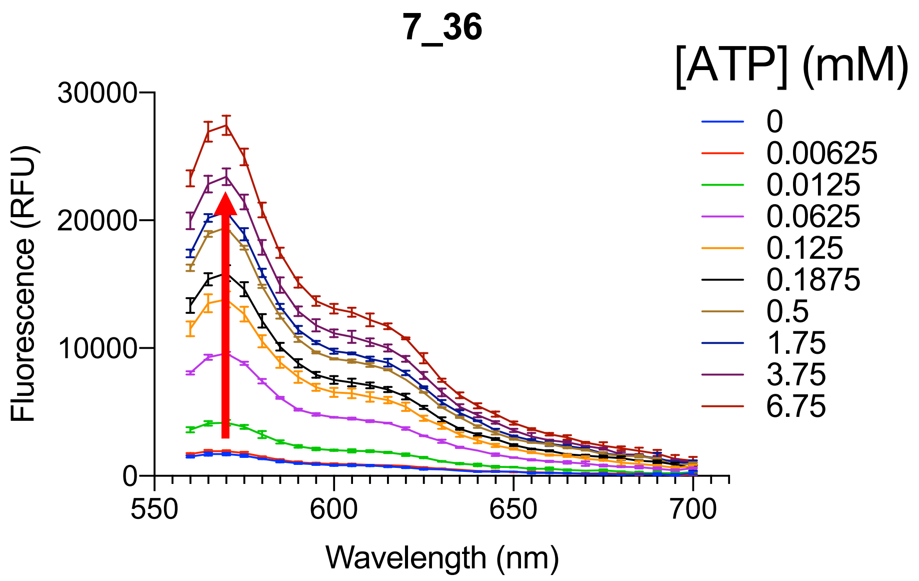


**b**

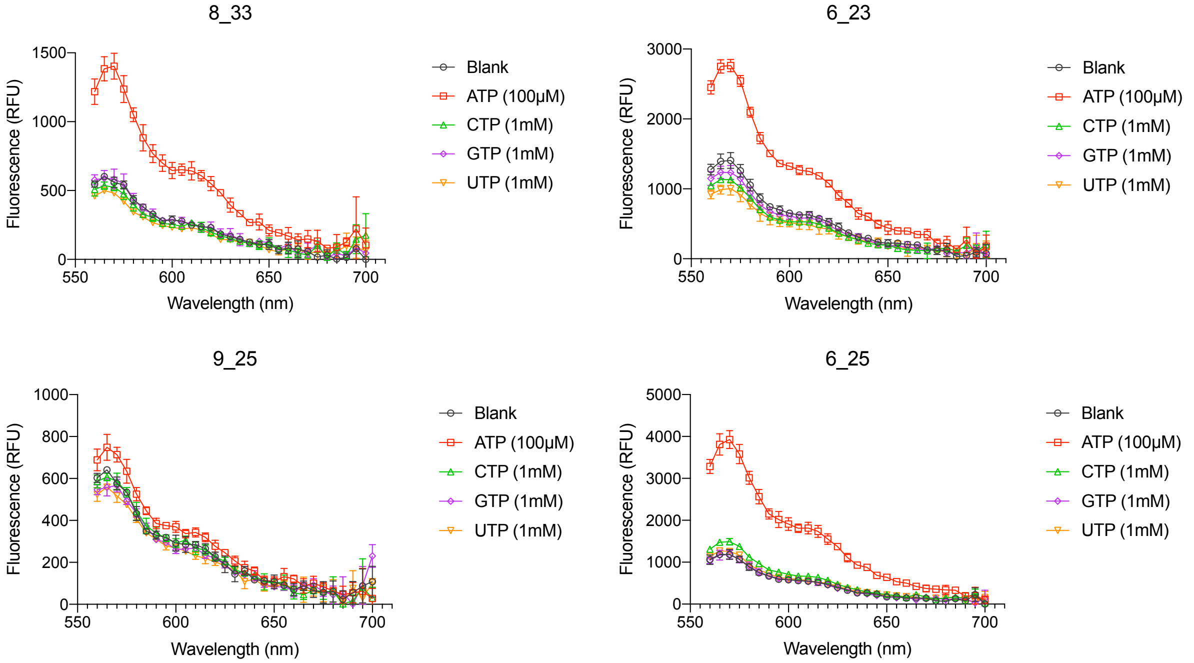


**Supplementary Figure 1 I Raw signal analysis and selectivity of constructs. (a)** Representative concentration-dependent emission spectra of an ISD switch. The fluorescence at peak emission (570 nm) was used as raw signal for both thermodynamic and kinetic studies. Concentrations of ATP in mM are listed. **(b)** Constructs with linker and double-stranded regions of varying lengths retain the high selectivity of the native aptamer for 100 μM ATP versus a 10-fold higher concentration of non-target ribonucleotides. All plots are averaged over n=3 replicates. Error bars represent the standard deviation.

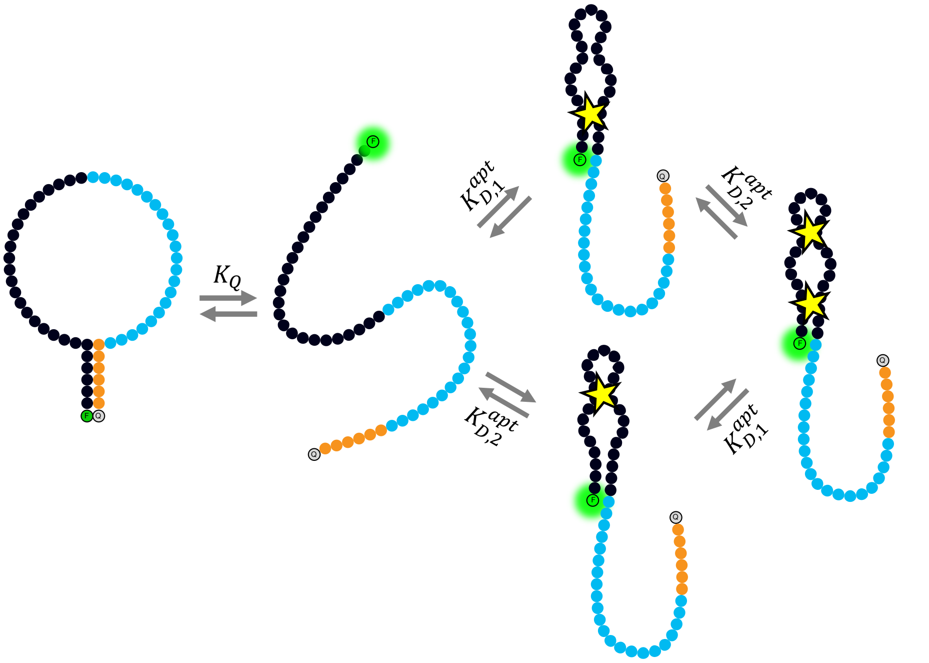


**Supplementary Figure 2 I Two-site binding model.** Here we derive the equations governing a two-site binding ISD construct. This model is specific to the ATP aptamer[1,2] and is likely to not be generalizable to other aptamers.

We recover the bimodal $K_D$ behavior by allowing site 1 and site 2 to vary in fluorescence values. Without this assumption, the above model would result in a binding curve with a concave second derivative. We assume binding to site 1 and binding to site 2 have different fluorescence values, $\eta_1$ and $\eta_2$. We extract $K_{D,1}$ and $K_Q$ via:

$$Signal = B_{max} \frac{K_{D,1}K_{D,2}+\eta_1 K_{D,1}[T]+\eta_2 K_{D,2}[T]+K_{D,1}K_{D,2}[T]^2}{K_{D,1}K_{D,2}(1+K_Q)+K_{D,1}[T]+K_{D,2}[T]+K_{D,1}K_{D,2}[T]^2} \qquad \text{(S1)}$$

In the manuscript, we report $K_D^{eff}$ as calculated by equation 4 using $K_{D,1}$.

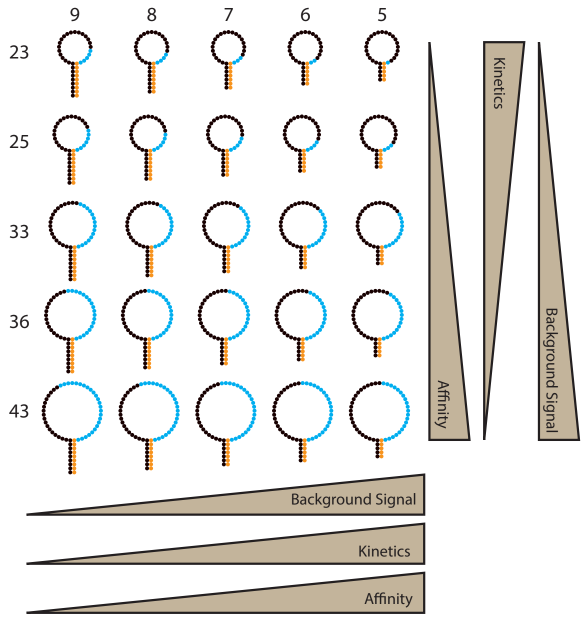


| | | $K_D^{eff}$ (mM) | | | $k_{obs}^{fast}$ ($s^{-1}$) | | |
|---|---|---|---|---|---|---|---|
| $L_{DS}$ | $L_{loop}$ | Average | | Std. Err. | Average | Upper CI | Lower CI |
| 9 | 23 | 39.0 | ± | 35.7 | 2.29 | 2.86 | 1.71 |
| | 25 | 12.8 | ± | 1.7 | 1.70 | 1.94 | 1.45 |
| | 33 | 4.99 | ± | 0.53 | 0.32 | 0.33 | 0.31 |
| | 36 | 2.72 | ± | 1.47 | 0.31 | 0.32 | 0.29 |
| | 43 | 0.968 | ± | 0.085 | 0.27 | 0.30 | 0.25 |
| 8 | 23 | 3.58 | ± | 1.38 | 3.58 | N/A | N/A |
| | 25 | 2.29 | ± | 0.05 | 3.61 | 4.02 | 3.21 |
| | 33 | 0.735 | ± | 0.01 | 1.51 | 1.56 | 1.47 |
| | 36 | 0.352 | ± | 0.043 | 1.77 | 1.85 | 1.69 |
| | 43 | 0.103 | ± | 0.114 | 1.43 | 1.48 | 1.39 |
| 7 | 23 | 0.382 | ± | 0.034 | 4.30 | 4.99 | 3.61 |
| | 25 | 0.272 | ± | 0.038 | 4.02 | 4.43 | 3.60 |
| | 33 | 0.0985 | ± | 0.0481 | 3.21 | 3.39 | 3.04 |
| | 36 | 0.0709 | ± | 0.0018 | 2.95 | 3.18 | 2.73 |
| | 43 | 0.0298 | ± | 0.0032 | 2.73 | 2.90 | 2.56 |
| 6 | 23 | 0.0314 | ± | 0.0111 | 5.82 | 9.17 | 2.47 |
| | 25 | 0.0265 | ± | 0.0027 | 4.18 | 4.71 | 3.65 |
| | 33 | 0.0182 | ± | 0.0076 | 4.53 | 5.25 | 3.82 |
| | 36 | 0.0107 | ± | 0.0019 | 4.82 | 5.65 | 3.99 |
| | 43 | 0.0149 | ± | 0.0122 | 3.31 | 3.72 | 2.90 |
| 5 | 23 | 21.4 | ± | 51.3 | N/A | N/A | N/A |
| | 25 | 0.0379 | ± | 0.0191 | N/A | N/A | N/A |
| | 33 | 9.2 | ± | 6.35 | N/A | N/A | N/A |
| | 36 | 0.0195 | ± | 0.0538 | N/A | N/A | N/A |
| | 43 | 2.85 | ± | 16.08 | N/A | N/A | N/A |

**Supplementary Figure 3 I Overview of results for various ISD constructs used in this work.** The constructs tested are graphically summarized (left). The table (right) summarizes all measured thermodynamic and kinetic parameters for these constructs. For constructs where $L_{DS}$ = 5, kinetics were faster than the time resolution of our detector, and we could not obtain a robust fit.

**Supplementary Figure 4 I Conformational selection model.** It has been suggested that the ATP aptamer used in this work can undergo both induced-fit[3] and conformational selection[4] binding mechanisms. Therefore, we present the equations governing ISD through a single-site, conformational selection binding mechanism. Since the pre-folded, binding-competent form also gives a signal, aptamers exhibiting this binding mechanism are more likely to suffer from larger background signal or possibly slower kinetics[5].

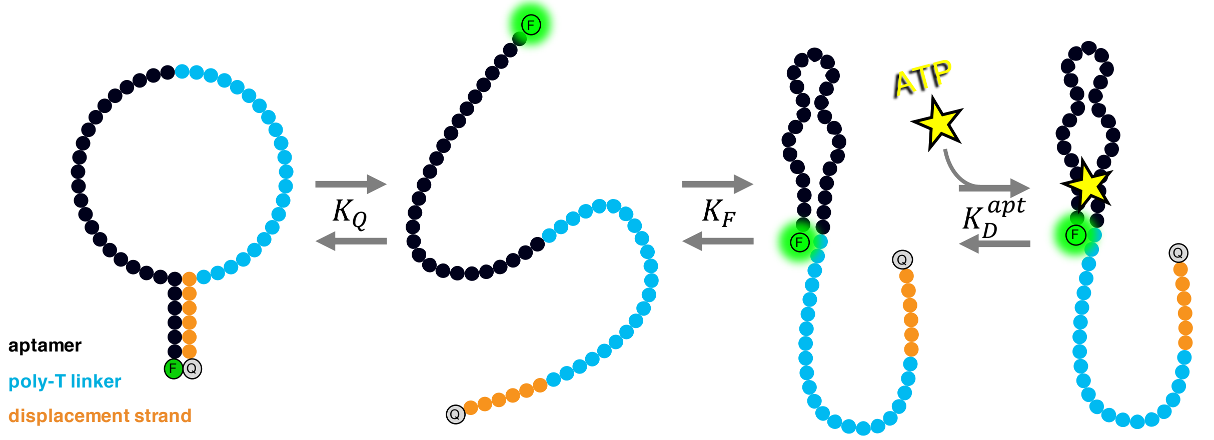


$$Signal = \frac{1+K_F+\frac{K_F[T]}{K_D^{apt}}}{K_Q+1+K_F+\frac{K_F[T]}{K_D^{apt}}} \quad \text{(S2)}$$

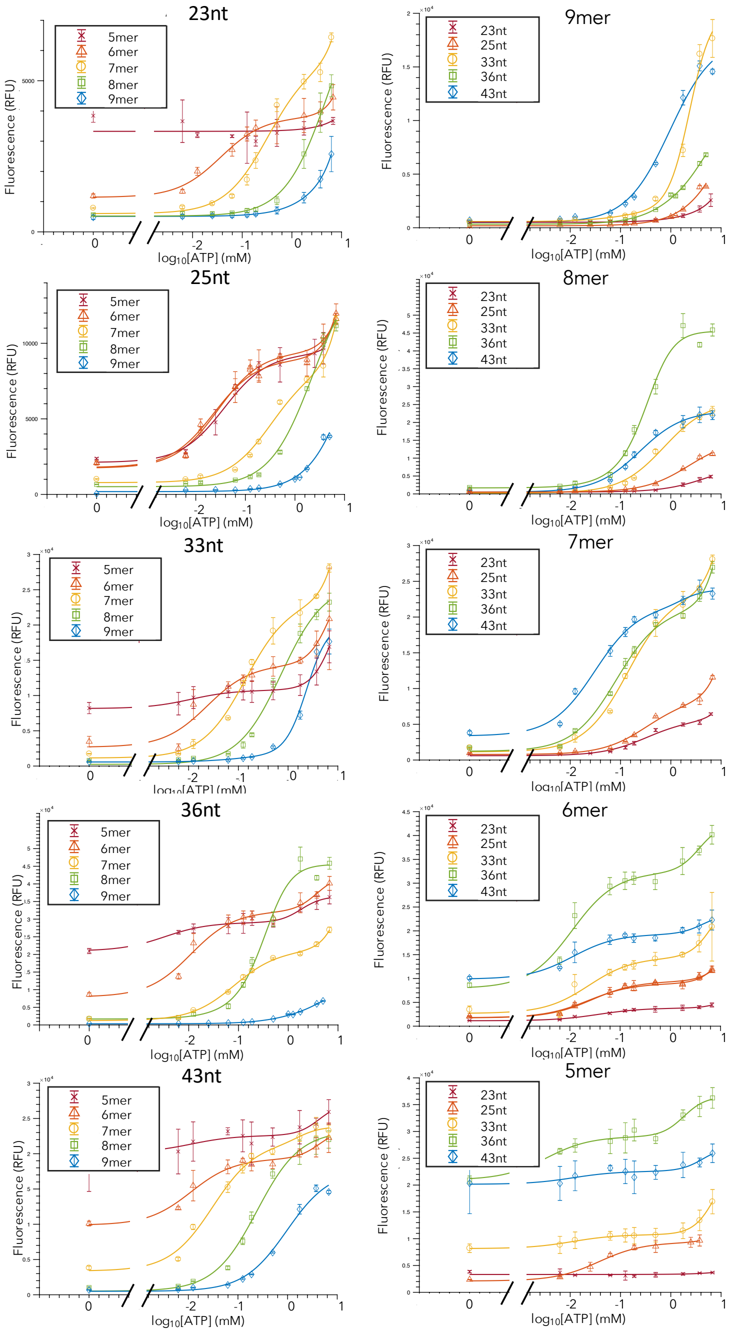


**Supplementary Figure 5 I Binding curves for all combinations of displacement strand and loop lengths.**

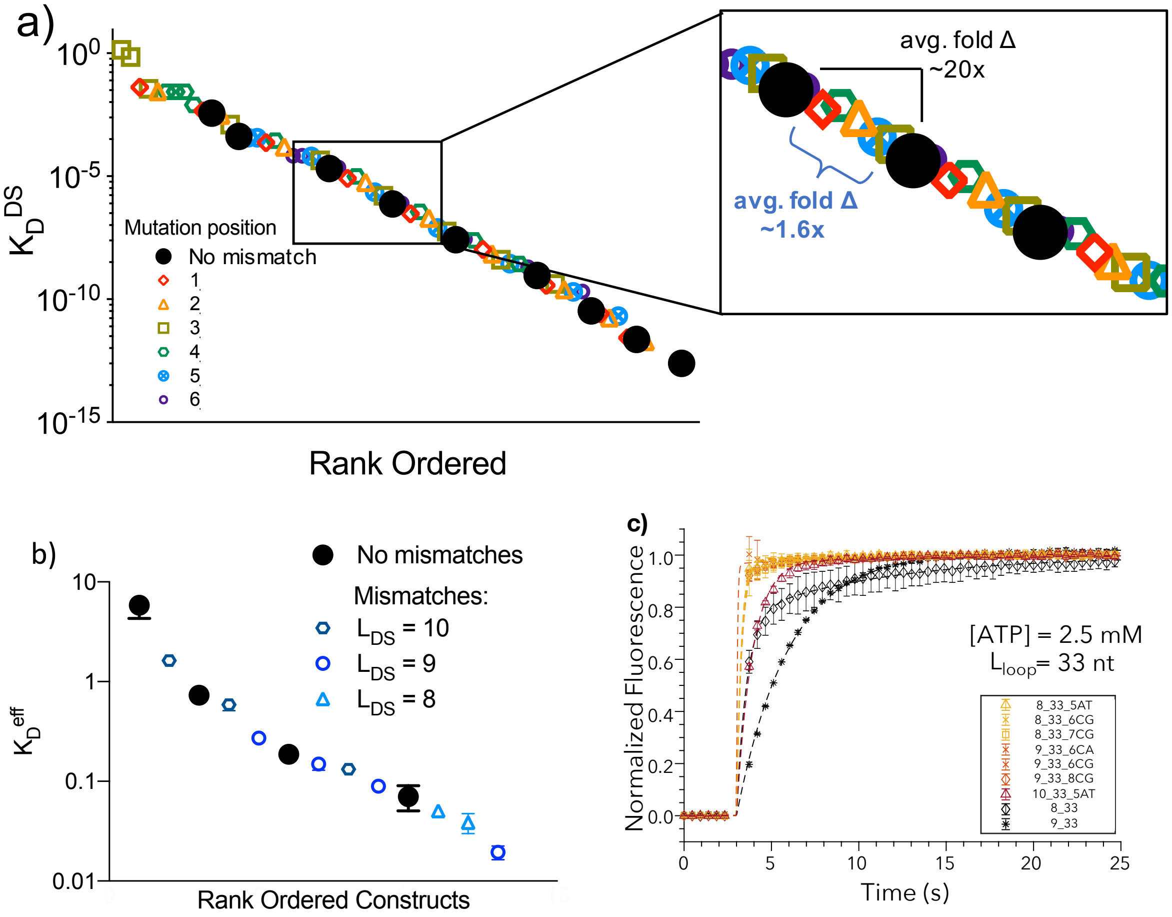


**Supplementary Figure 6 I Effects of mismatch incorporation.** (**a**) When considering perfectly matched, unlinked displacement strands (black circles) the set of theoretically obtainable $K_D^{DS}$ features very large jumps (20.6 ± 8.7 per base). On the other hand, incorporating single mismatches (colored symbols) greatly decreases this spacing to 1.6 ± 0.8. Calculations were performed using $\Delta G_{fold}$ from *mfold* at room temperature and 6 mM $Mg^{2+}$. Mutation position is defined from the 3'-end of the displacement strand. (**b**) Measured effective binding affinities for perfectly matched strands (black circles) and mismatches (colored symbols) with $L_{loop}$ = 33 nt. Adding mismatches greatly increases the tunability of the thermodynamics. (**c**) Introducing mismatches greatly increases the signaling kinetics of the constructs, e.g. yellow vs. black star and orange vs. black diamond. Even a 10mer DS with mismatches (red) exhibits kinetics faster than that of the corresponding aptamer beacon[6].

**Supplementary Calculation 1 I**

The relative impacts of $L_{loop}$ and $L_{DS}$ on $K_Q$ can be calculated as follows:

$$K_Q \sim \frac{\exp\left(\frac{1.7 L_{DS}}{RT}\right)}{(L_{loop})^{2.6}} \quad \text{(S3)}$$

$$\frac{dK_Q}{dL_{DS}} \sim \frac{1.7}{RT} K_Q \quad \text{(S4)}$$

$$\frac{dK_Q}{dL_{loop}} \sim -\frac{2.6}{L_{loop}} K_Q \quad \text{(S5)}$$

At room temperature, we find:

$$\frac{dK_Q}{dL_{loop}} \sim \frac{-7.5}{L_{loop}} \frac{dK_Q}{dL_{DS}} \quad \text{(S6)}$$

Fits to **Figure 3b** and **d** yield $L_{loop} \frac{\frac{dK_Q}{dL_{loop}}}{\frac{dK_Q}{dL_{DS}}} = 6.0 \pm 3.4$, which is in agreement with our theory.

More specifically, linear fits to **Figure 3b** and **d** yield:

$$a = \left.\frac{d\log K_Q}{dL_{loop}}\right|_{constant\ L_{DS}} = -0.049 \pm 0.030 \ \frac{log\ change\ K_Q}{loop\ nt}$$

$$b = \left.\frac{d\log K_Q}{dL_{DS}}\right|_{constant\ L_{loop}} = 0.827 \pm 0.154 \ \frac{log\ change\ K_Q}{DS\ bp}$$

Where $\sigma$ = standard deviation over the five combinations of linker or DS.

The equivalence between loop bases and displacement strand bases is then given by:

$$\frac{n_{loop}}{n_{DS}} = \frac{\left.\frac{d\log K_Q}{dL_{DS}}\right|_{constant\ L_{loop}}}{\left.\frac{d\log K_Q}{dL_{loop}}\right|_{constant\ L_{DS}}} = -17.7 \pm\ 11.9$$

Where $\sigma$ is calculated using propagation of errors: $\sigma = \sqrt{\left(\frac{\sigma_b}{a}\right)^2 + \left(\frac{\text{b}\ \sigma_a}{a^2}\right)^2}$

The loop/DS equivalence varies over $L_{DS}$ and $L_{loop}$ and is summarized below:

| $\frac{n_{loop}}{n_{DS}}$ | 23 | 25 | 33 | 36 | 43 |
|---|---|---|---|---|---|
| 5 | -95.2 ± 551.2 | -83.4 ± 483.0 | -76.0 ± 439.9 | -73.4 ± 425.0 | -56.2 ± 325.4 |
| 6 | -53.4 ± 19.7 | -46.8 ± 17.1 | -42.6 ± 15.7 | -41.2 ± 15.1 | -31.5 ± 12.2 |
| 7 | -18.7 ± 1.5 | -16.4 ± 1.0 | -15.0 ± 1.1 | -14.4 ± 1.0 | -11.1 ± 1.6 |
| 8 | -13.6 ± 1.3 | -11.9 ± 1.0 | -10.9 ± 1.0 | -10.5 ± 1.0 | -8.0 ± 1.2 |
| 9 | -13.9 ± 1.6 | -12.2 ± 1.3 | -11.1 ± 1.2 | -10.7 ± 1.2 | -8.2 ± 1.3 |

**Supplementary Calculation 2 I Derivation of the signaling kinetics of an induced fit binding mechanism.** Kinetics for induced fit can be derived as follows (see Mathematica code at https://github.com/btotherad77/isd for full analytical solution). Importantly, the analysis reveals two effective time constants—a fast time constant and a slow time constant—that are functions of the parameters of both aptamer and displacement strand kinetics. Since we assume the kinetics of aptamer binding are fixed, we obtain control of $k_{obs}$ by modulating $k_{on}^{DS}$ and $k_{off}^{DS}$.

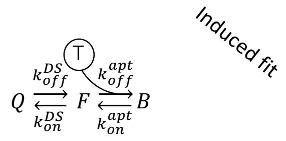


$$\underline{C(t)} = \begin{pmatrix} C_Q(t) \\ C_F(t) \\ C_B(t) \end{pmatrix}$$

$$\underline{\underline{k}} = \begin{pmatrix} -k_{off}^{DS} & k_{on}^{DS} & 0 \\ k_{off}^{DS} & -(k_{on}^{DS} + k_{on}^{apt}[T]) & k_{off}^{apt} \\ 0 & k_{on}^{apt}[T] & -k_{off}^{apt} \end{pmatrix}$$

$$\frac{d}{dt}\underline{C(t)} = \underline{\underline{k}} \bullet \underline{C(t)}$$

$$\underline{C(t)} = \exp\left(\underline{\underline{k}}t\right) \bullet \underline{C_0}$$

$$\underline{C(t)} = \underline{\underline{x}}\exp\left(\underline{\underline{\Lambda}}t\right)\underline{\underline{x}}^{-1} \bullet \underline{C_0}$$

eigenvectors of *k*

$$\exp\left(\underline{\underline{\Lambda}}t\right) = \begin{pmatrix} \exp(\lambda_0 t) & \cdots & 0 \\ \vdots & \ddots & \vdots \\ 0 & \cdots & \exp(\lambda_n t) \end{pmatrix}$$

eigenvalues

Analytical solution reveals two time constants:

$$= \frac{1}{2}\left(k_{off}^{DS} + k_{on}^{DS} + [T]k_{on}^{apt} + k_{off}^{apt} + \sqrt{\left(k_{off}^{DS} + k_{on}^{DS} + [T]k_{on}^{apt} + k_{off}^{apt}\right)^2 - 4\left(k_{on}^{DS}k_{off}^{apt} + k_{off}^{DS}\left(k_{off}^{apt} + [T]k_{on}^{apt}\right)\right)}\right)$$

$$= \frac{1}{2}\left(k_{off}^{DS} + k_{on}^{DS} + [T]k_{on}^{apt} + k_{off}^{apt} - \sqrt{\left(k_{off}^{DS} + k_{on}^{DS} + [T]k_{on}^{apt} + k_{off}^{apt}\right)^2 - 4\left(k_{on}^{DS}k_{off}^{apt} + k_{off}^{DS}\left(k_{off}^{apt} + [T]k_{on}^{apt}\right)\right)}\right)$$
